# Optimized R2 Retroelement Complexes Enable Precise and Efficient DNA Insertion into Plant Genomes

**DOI:** 10.1101/2025.08.22.671877

**Authors:** Kimberley T. Muchenje, Yunqing Wang, Tufan M. Oz, Eugene Li, Amelia Saffron, Gozde S. Demirer

## Abstract

Precise, targeted insertion of multi-kilobase DNA sequences into plant genomes is critical for studying gene function, ensuring robust transgene expression, and stacking traits in crops, but remains challenging. Existing targeted insertion methods in plants relying on programmable nucleases are inefficient and can generate unwanted mutations. Newer technologies based on prime editors, transposases, and site-specific recombinases extend capabilities but remain constrained with low efficiencies, off-target integration, silencing, or limited DNA payload size. R2 non-long terminal repeat (non-LTR) retrotransposons integrate via target-primed reverse transcription (TPRT) specifically targeting the 25S ribosomal DNA multicopy site and enabling double-strand-break-free installation of gene-sized DNA sequences. We adapted the avian *Taeniopygia guttata* R2 protein (R2Tg) for targeted DNA insertion into plant genomes through engineering of R2Tg expression cassettes and RNA payloads carrying intron-disrupted mCherry and RUBY retrotransposition reporters with length-optimized rDNA homology arms. These efforts, together with optimized construct delivery formats and incubation temperatures, define R2 editor design rules enabling efficient DNA integration and functional protein expression from the 25S rDNA locus. In *Nicotiana benthamiana* leaves, *Arabidopsis thaliana* protoplasts, and *Solanum lycopersicum* seedlings, the optimized R2Tg editor system achieved targeted insertion with efficiencies up to 24% payloads ranging in size from 2kb to 5kb. This work establishes a compact R2Tg ribonucleoprotein platform for targeted DNA insertion into plant genomes, targeting a multicopy genomic safe harbor site to enable efficient multi-kilobase gene addition

## Introduction

Crop yields are increasingly threatened by rising global temperatures, intensifying drought, and the expanding range of plant diseases^1^. Genome engineering offers a powerful, timely strategy for enhancing climate resilience, food security, and sustainable biomanufacturing^2,3^. The development of improved crops relies heavily on the incorporation of foreign genes and heterologous metabolic pathways into plant genomes, where most approaches use *Agrobacterium tumefaciens* for DNA delivery and insertion. However, the random DNA integration of *Agrobacterium* can often cause variable gene expression due to local chromatin context and nearby regulatory elements, lead to silencing, and prevent precise tagging of specific genes or loci, all of which are limitations complicating functional genomics studies and requiring extensive screening across generations to identify lines with desired, stable, and heritable expression^4–7^.

Emerging targeted insertion technologies allow controlled placement of DNA sequences at predetermined genomic loci, eliminating risks of unintended gene expression changes and genomic instability^8^. Furthermore, precise DNA insertion at defined genomic sites is essential for probing gene function and regulation and enables robust outputs in synthetic gene circuits, efficient trait stacking, and accelerated development of engineered crops for agriculture and bioproduction^9,10^. To date, multiple strategies have been developed to achieve site-specific DNA integration in plants. Programmable nucleases, such as CRISPR-Cas9, introduce targeted double-strand breaks (DSBs), which are repaired by host machinery^11,12^. Although homology-directed repair can result in precise integration using donor templates with flanking homology arms at the DSB site^13^, this pathway is not preferred in most plant cells, where DNA breaks are predominantly repaired by non-homologous end joining, often leading to low efficiency, imprecise insertions, and indel generation^14^. Moreover, DSBs can lead to unpredictable outcomes, such as large deletions or chromosomal rearrangements^15^.

As an alternative, prime editing offers a DSB-free approach for introducing precise nucleotide changes and short sequence insertions into the genome^16,17^. Despite broad applicability, its efficiency to insert large DNA segments in plants remains limited, restricting its use for integrating full-length genes or metabolic pathways^18^. Expanding the capabilities of prime editing, PrimeRoot employs optimized prime editing guide RNA (pegRNA) architectures, prime editors, and site-specific recombinases to achieve DSB-free targeted insertion of large DNA sequences into plant genomes^19^. Although promising with up to 6.3% efficiency, its broader application may be limited by lack of generalizable pegRNA design rules to ensure robust activity across diverse loci, and the low efficiency of the recombinase recognition site installation, which is a critical first step in PrimeRoot. These challenges highlight the need for continued engineering of more effective editors to fully realize the potential of precise, programmable DNA insertion in plants.

Expanding the toolkit further, transposon-based systems offer another alternative for programmable insertion. Transposons are mobile elements that can be broadly categorized as DNA transposons or retrotransposons based on their integration intermediates. They possess sophisticated insertion mechanisms setting them up as powerful tools for targeted insertion^20,21^. A recent discovery of CRISPR-associated DNA transposases (CAST) in bacterial genomes enabled the development of editors that use programmable guide RNAs to direct transposon insertion to defined genomic sites^22,23^. Unlike nuclease-based editors, they lack DNA cleavage activity and instead rely entirely on transposase machinery to perform the integration. Even though their translation to eukaryotic human cells has been successful – albeit with a low efficiency^24^ – native bacterial CAST system has yet to demonstrate feasibility in plants. However, building on this discovery, transposase-assisted target-site integration (TATSI) system was developed, which co-expresses Cas9 nuclease with the rice Pong DNA transposase to enable high efficiency, targeted DNA insertion in *Arabidopsis thaliana* and soybean^25^. While TATSI offers a promising approach for targeted DNA insertion in plants, its performance can be affected by high rates of off-target integration at native transposase recognition motifs, reliance on DSBs, and increased susceptibility to silencing due to the use of plant-derived transposases.

In addition to DNA transposons, retrotransposons have also shown recent promise for targeted gene insertion in eukaryotes. R2 non-long terminal repeat (LTR) retrotransposons are a class of sequence-specific mobile elements that insert into the conserved 28S ribosomal DNA (rDNA) tandem repeat loci and have so far been identified in the genomes of multicellular animals including insects, crustaceans, and non-mammalian vertebrates^26–28^. R2 proteins have been successfully repurposed as high-efficiency genome editors in mammalian systems, both at their conserved target sites and at other RNA-guided genomic loci through Cas9 fusion, leveraging their target-primed reverse transcription (TPRT) mechanism^29–32^. Engineered R2 retrotransposons mediate RNA-templated insertions at the native 28S rDNA locus with efficiencies up to 80% in HEK293T cells and 60% in mouse embryos, and they achieve up to 40% at Cas9-directed, non-rDNA sites^30–32^.

R2 retrotransposons provide a unique, DSB-free approach for targeted DNA insertion into the 28S rDNA site, which is referred to as 25S rDNA site in plants, offering broad adaptability for genome editing across plant species. The required domains for function, DNA binding, endonuclease, and reverse transcriptase, are encoded within a single open reading frame, enabling a compact editor design that can be co-delivered with its RNA payload as a ribonucleoprotein (RNP) complex. Moreover, the high copy number of rDNA repeats in plants (∼800 copies in *A. thaliana*^33^, ∼1247 copies in *Nicotiana benthamiana*^34^, ∼1200 copies in *Solanum lycopersicum*^35,36^ and ∼6650 copies in wheat^37^) can further increase integration frequency and expression, and simplify downstream screening for precise insertion events.

Here, we established and optimized a plant genome editor based on the R2 retrotransposon that enables targeted, high-efficiency DNA insertion into the 25S rDNA loci. This system combines: i) an optimized R2 expression, ii) RNA donor carrying mCherry and RUBY retrotransposition reporters with length-optimized rDNA-homology arms, and iii) systematically tuned 5′ and 3′ UTRs in the RNA template. The optimized system achieves integration of payloads up to 5kb size in *Nicotiana benthamiana* leaves, *Arabidopsis thaliana* protoplasts, and *Solanum lycopersicum* seedlings with efficiencies reaching 24% of transformed cells quantified by imaging and 0.9-1 copies per genome quantified by droplet digital PCR on both 5′ and 3′ integrated junctions. R2-mediated insertion events were validated at the 25S rDNA loci with molecular evidence confirming both high efficiency and target specificity, establishing a robust platform for precise DNA insertion into plant genomic safe-harbor loci for various fundamental plant biology or biotechnology applications.

## Results

### Establishing an R2 retrotransposon-mediated platform for targeted DNA insertion in plants

Building on the well-characterized target-primed reverse transcription (TPRT) activity of R2 proteins in mammalian cells^27,28^, we investigated whether the same machinery could mediate precise DNA insertion in the plant nuclear genome. In the canonical mechanism, an R2 ribonucleoprotein complex composed of an R2 protein and its RNA template binds the 28S rDNA locus (25S rDNA in plants) in a manner that depends on the secondary structures of RNA sequences within the 5′ and 3′ untranslated regions (UTRs)^27,28^. The complex then nicks the bottom strand, exposing a 3′-hydroxyl primer, and synthesizes first-strand complementary DNA (cDNA) from its RNA template^27^. Subsequently, the R2 protein nicks the top strand, and initiates second-strand synthesis^28^, where the integrated element is flanked by the R2 5′ and 3′ UTRs (**Fig. 1A**).

**Figure 1.**
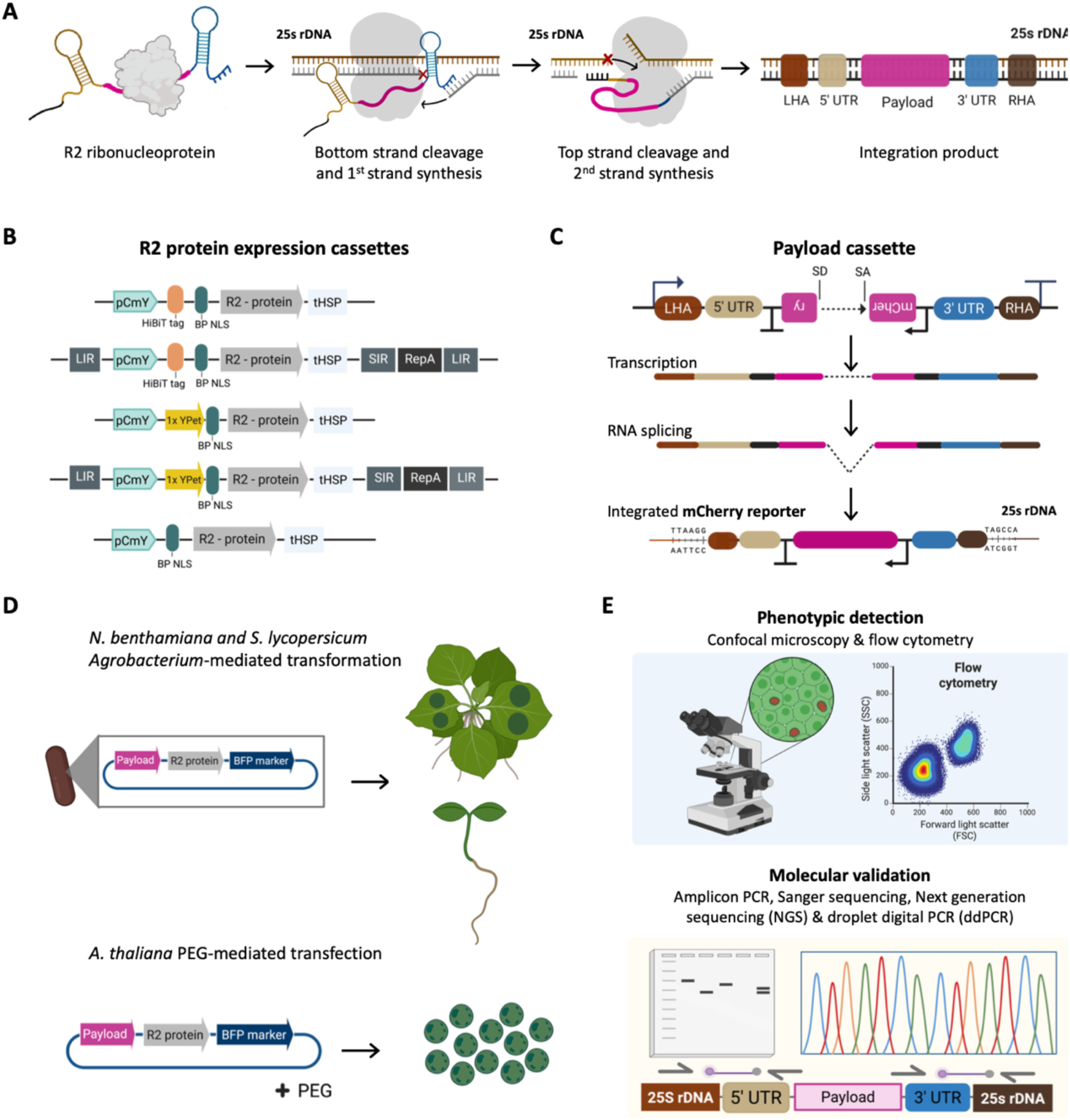
Overview of R2 retrotransposon-mediated targeted DNA insertion in plants. **(A)** Mechanism of R2 integration at the 25S rDNA locus via site-specific Target Primed Reverse Transcription. **(B)** Constructs for detecting R2 protein expression in plants, with geminiviral (GV) parts added for enhanced expression. Proteins were tagged at the N-terminus with HiBiT tag for luminescence detection or 1x YPEt for fluorescence detection. All proteins were also tagged at the N-terminus with bipartite (BP) NLS tag for nuclear localization. **(C)** Payload design: An mCherry intron-containing cassette embedded within 5′ and 3′ UTRs is spliced to generate a mature RNA template for retrotransposition, enabling phenotypic detection of integration by confocal microscopy. **(D)** R2 system expression was performed via *Agrobacterium tumefaciens* infiltration of *Nicotiana benthamiana* leaves and *Solanum lycopersicum* seedlings as well as PEG-mediated *Arabidopsis thaliana* protoplast transfection. **(E)** Integration events were evaluated by microscopy, flow cytometry, amplicon PCRs, NGS, and droplet digital PCR (ddPCR). HiBiT: high affinity bioluminescence tag, YPet: yellow fluorescent protein, BFP: blue fluorescent protein, BP NLS: Bipartite nuclear localization signal, UTR: untranslated region, rDNA: ribosomal DNA, LHA: left homology arm, RHA: right homology arm, pCmY: pCmYLCV6 promoter derived from Cestrum yellow leaf-curling virus, tHSP: *Arabidopsis thaliana* heat shock protein 18.2 terminator, GV: geminiviral replicon, LIR: long intergenic region of the geminiviral replicon, SIR: short intergenic region of the geminiviral replicon, RepA: trans-acting replication-initiation protein. SD: splice donor, SA: splice acceptor.

We deployed the R2 retrotransposon system (R2 system in short) to leverage this double-strand-break-free integration mechanism, combined with the high copy number^34,37^ and transcriptional activity^38,39^ of the 25S rDNA array, to achieve robust expression of integrated gene sequences in plants. Efficient TPRT requires high amounts of nuclear-localized R2 protein^40^. We therefore codon-optimized R2 open reading frames for plant expression, appended an N-terminal strong bipartite nuclear localization signal (BP NLS), and fused either a high affinity bioluminescence tag (HiBiT) for luminescence complementation or a single-copy YPet fluorescent protein to the N-terminus for quantifying and/or imaging protein expression (**Fig. 1B**). Each construct was driven by the strong pCmYLCV6 promoter and terminated by the *A. thaliana* tHSP 18.2 terminator. To further increase intracellular R2 copy number, long and short intergenic regions (LIR-SIR) together with the RepA replication-initiation protein were incorporated as a second set to test, generating self-replicating geminiviral (GV) replicons that boost transient expression^41^ (**Fig. 1B**).

To enable simple phenotypic detection of R2-mediated insertion events, we built an RNA payload to function as a retrotransposition reporter following a classic GFP reporter^42,43^, in which either the *Solanum tuberosum* ST-LS1 intron^44^ or an *Arabidopsis thaliana* intron^45^-disrupted mCherry cassette was embedded between the 5′ and 3′ untranslated regions (UTRs) of the R2 element (**Fig. 1C**). Following transcription and intron splicing, the resulting RNA forms a contiguous template for reverse transcription via R2 protein. Hence, precise integration via R2 system restores an intron-less mCherry expression cassette, producing a fluorescent signal that enables visualization and quantification by confocal microscopy.

To evaluate R2 system activity across different contexts, we introduced R2 protein and RNA payload constructs into *Nicotiana benthamiana* leaves and *Solanum lycopersicum* seedlings via *Agrobacterium tumefaciens* infiltration, and into *A. thaliana* protoplasts using polyethylene glycol (PEG)-mediated transfection (**Fig. 1D**). BFP fluorescent marker was included in the constructs for efficiency normalization. Phenotypic detection was achieved via confocal microscopy for *N. benthamiana* and *S. lycopersicum* leaves and flow cytometry for *A. thaliana* protoplasts. Molecular characterization of insertion junctions and copy number was performed using amplicon PCR, Sanger sequencing, high-depth next-generation sequencing (NGS), and droplet digital PCR (ddPCR) (**Fig. 1E**).

### R2 expression optimization and functional screening identify an active R2 system in plants

To identify a functional R2 editor in plants, we first compared the expression profiles of four codon-optimized R2 proteins: R2Bm from *Bombyx mori*, R2Za from *Zonotrichia albicollis*, R2Tg from *Taeniopygia guttata*, and a rationally improved variant of R2Tg, R2Tg-OPT^30^. These proteins were expressed in *N. benthamiana* leaves using Agro-infiltration. All proteins were tested with and without geminiviral (GV) replicon elements to evaluate the impact of expression cassette copy number on R2 protein accumulation (**Fig. 1B**). 3 days after infiltration, we detected the expression of all YPet-tagged R2 proteins using confocal microscopy, where nuclear-localized YPet signal was consistently stronger in GV versions (**Fig. 2A**). Luminescence quantification using a plate reader for the HiBiT-tagged set of proteins in the plant nuclear fraction confirmed that GV parts significantly increased R2 protein levels across all variants (**Fig. 2B**). These results established our ability to express high levels of all tested R2 proteins in plants cells and that GV replicons are an effective strategy for increasing R2 protein levels in the nuclei of *N. benthamiana* leaf cells.

**Figure 2.**
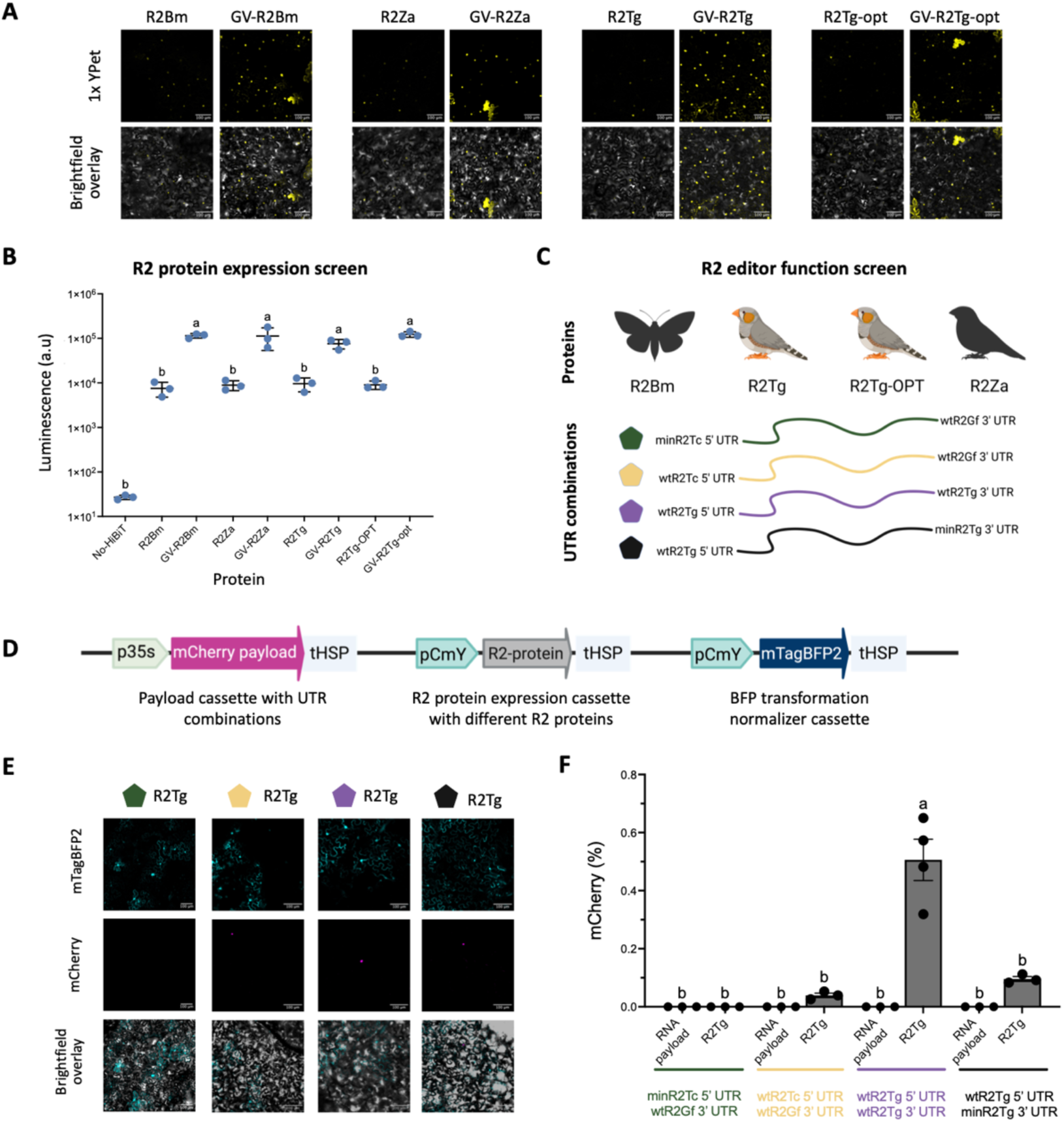
Protein expression and function screening identify optimal R2 proteins and UTR variant combinations. **(A)** Representative confocal fluorescence microscopy images of *N. benthamiana* leaves expressing 1×YPet-tagged R2 proteins with or without geminiviral (GV) replication elements. Scale bars, 100 µm. **(B)** R2 protein expression quantified by HiBiT luminescence (n=3 biological replicates). Ordinary one-way ANOVA Tukey HSD multiple comparison test. Different letters denote statistically significant differences. Values are mean ± S.E.M. See Source Data File for details. **(C)** R2 editor screen setup with combinations of four UTRs and four R2 proteins. **(D)** All-in-one screening construct containing an mCherry payload cassette, R2 protein cassette, and an mTagBFP2 transformation marker. **(E)** Representative confocal microscopy images of *N. benthamiana* leaves showing mTagBFP2 (cyan) indicating transfection and mCherry (magenta) indicating R2-mediated targeted insertion. Scale bars, 100 µm. **(F)** R2-mediated targeted insertion efficiency quantified from confocal microscopy images (n=3 biological replicates). Ordinary one-way Anova with Tukey HSD multiple comparison test. Different letters denote statistically significant differences. Values are mean ± S.E.M. See Source Data File for details. UTR: untranslated region, min: minimal sequence variant, wt: wild-type sequence, p35S: Cauliflower mosaic virus 35S promoter, pCmY: pCmYLCV6 promoter, tHSP: heat shock protein 18.2 terminator, mTagBFP2: blue fluorescent protein, mCherry: red fluorescent protein, a.u: arbitrary units.

Previous mammalian studies have shown that autocatalytic cleavage of the 5′ UTR is required to generate competent RNA templates for R2-mediated integration via TPRT^30^. To assess whether the 5′ UTR sequences used in our donor constructs enable autocleavage, we performed *in vitro* transcription and cleavage assays using the wild-type and minimal length *T. castaneum* and *T. guttata* 5’ UTR, each fused to 100 bp of the rDNA sequence upstream of the insertion site, referred to here as the left homology arm (LHA) (**Supplementary Fig. 1A**). Distinct cleavage products were observed by agarose gel electrophoresis at expected band sizes, confirming that all tested UTRs retain autocatalytic activity *in vitro* (**Supplementary Fig. 1B**).

Next, we performed a systematic R2 editor function screen to identify an optimal pairing of R2 protein and donor RNA architecture. Each configuration combined one of the four R2 proteins with a donor cassette encoding the mCherry-based retrotransposition reporter and different UTR combinations (**Fig. 2C**). We evaluated four UTR combinations, all previously shown to be active in mammalian systems, derived from three species: minimal *T. castaneum* 5′ UTR with wild-type *Geospiza fortis* 3′ UTR; wild-type *T. castaneum* 5′ UTR with wild-type *G. fortis* 3′ UTR; wild-type *T. guttata* 5′ and 3′ UTRs; and wild-type *T. guttata* 5′ UTR paired with a minimal *T. guttata* 3′ UTR^29,30^ (**Fig. 2C**), generating a library of 16 R2-UTR combinations to assess. Each all-in-one construct for the R2 editor function screen also encoded a constitutive mTagBFP2 reporter to normalize for transformation efficiency (**Fig. 2D**).

To evaluate functional activity, 7 days after infiltration, we imaged *N. benthamiana* leaves infiltrated with constructs expressing all-in-one constructs for each of the 16 R2 protein-UTR combinations and scored mCherry fluorescence as a direct readout of successful retrotransposition into the 25S rDNA (**Fig. 2E**). R2Bm and R2Za proteins did not yield detectable integration activity when paired with any of 4 UTR combinations in our editor screen (**Supplementary Fig. 1C, D**). The highest integration efficiency, defined as the percentage of leaf cells with detectable mCherry signal, was obtained with wild-type R2Tg paired with its wild-type 5′ and 3′ UTRs at ∼0.5% efficiency (**Fig. 2E, F**). Two additional R2Tg editor configurations with different UTRs also yielded successful integration (**Fig. 2E, F; Supplementary Fig. 1D**): R2Tg enabled detectable integration when paired with wild-type 5′ and minimal 3′ UTRs from *T. guttata* (0.11%), and wild-type *T. castaneum* 5′ UTR and wild-type *G. fortis* 3′ UTR (0.05%) (**Fig. 2E, F**). In contrast, no mCherry signal was observed when R2Tg was paired with the minimal *T. castaneum* 5′ UTR and wild-type *G. fortis* 3′ UTR (**Fig. 2E, F**). R2Tg-OPT exhibited minimal activity only when paired with wild-type *T. guttata* UTRs, and none of the other UTR combinations supported detectable integration by this variant. Our negative control samples consisting only of the payload without R2 protein did not generate any mCherry signal as expected (**Fig. 2F; Supplementary Fig. 2**)

This preliminary R2 editor function screen established the wild-type R2Tg protein, paired with its wild-type 5′ and 3′ UTRs, as the most efficient R2 editor configuration, showing potential for further optimization via R2 protein and payload engineering. We next focused on increasing the integration efficiency of the R2Tg editor paired with the mCherry payload containing its wild-type 5′ and 3′ UTRs with a payload length of 2.9kb, which we refer to as the R2Tg editor system.

### R2Tg system optimization via improved protein & payload expression and heat shock treatment

To ensure that mCherry fluorescence in our optimization assays only reflects bona fide R2-mediated integration events, we used an intron-disrupted mCherry retrotransposition reporter, same as above (**Fig. 3A**). This payload comprises an mCherry coding sequence in the antisense orientation relative to the RNA payload, interrupted by a plant intron in the sense orientation, flanked by R2TG wild-type 5′ and 3′ UTRs and 100 bp homology arms corresponding to the 25S rDNA target site. Transcription of the integrated cassette can only yield mCherry signal if the payload transcript is precisely spliced to remove the intron prior to R2-mediated insertion at the target locus. *Agrobacteria* T-DNA insertions or leaky expression cannot generate a functional, intron-free mCherry transcript because the cassette remains intron-bearing or incorrectly oriented, with the splice donor and acceptor motifs are present only on the sense strand, thereby ensuring the absence of false-positive signal in downstream fluorescence readouts.

**Figure 3.**
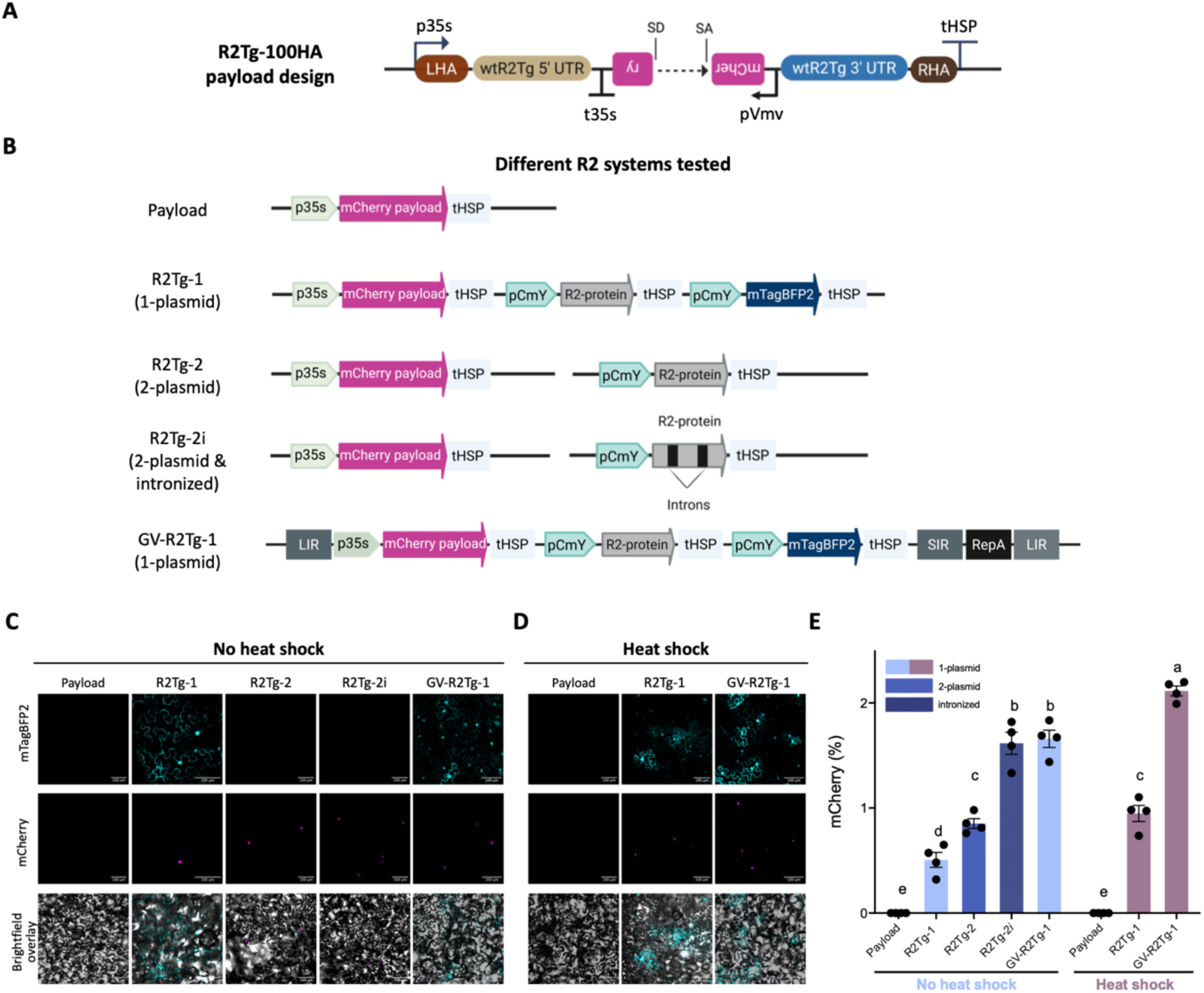
Higher R2 system copy number and heat shock treatment increases integration efficiency. **(A)** Schematic of the R2Tg payload cassette with 100 bp homology arms. **(B)** Schematic of the wild-type R2 editors using either 1-plasmid or 2-plasmid delivery approaches, intronized R2Tg-2i setup, and geminiviral-amplified R2 editor (GV-R2Tg-1). Representative confocal microscopy images of leaves infiltrated with R2 constructs **(C)** under no heat shock conditions and **(D)** heat shock; mTagBFP2 (cyan) is a transfection control; mCherry (magenta) reports successful R2-mediated integration at 25S rDNA loci. Scale bars, 100 µm. (**E)** Quantification of mCherry positive nuclei as a measure of targeted insertion efficiency. Bars represent mean ± S.E.M. of 4 independent biological replicates. P values were obtained using ordinary one-way ANOVA with Tukey HSD multiple comparison test. Different letters denote statistically significant differences. See Source Data File for details. LHA: left homology arm, RHA: right homology arm, p35S: Cauliflower mosaic virus 35S promoter, t35S: Cauliflower mosaic virus 35S terminator, pVmV: Cassava vein mosaic virus promoter, SD: splice donor, SA: splice acceptor, pCmY: pCmYLCV6 promoter, tHSP: heat shock protein 18.2 terminator, mTagBFP2: blue fluorescent protein variant, mCherry: red fluorescent protein, SIR: short intergenic region and LIR: long intergenic region of the geminiviral replicon, RepA: geminiviral replication-associated protein.

Using this reporter, we next tested whether targeted integration efficiency could be improved. Retrotransposon sequences are rapidly targeted by RNA-directed DNA methylation and converted to a stably silenced, heterochromatic state in plant cells^46–49^. We suspected that transcriptional interference^50^ from all cassettes contained in the all-in-one construct and silencing of R2 editor encoding sequences may be limiting the efficiencies in our initial R2 editor screen. To mitigate this effect, we split the system into two plasmids: one encoding the R2Tg protein and the other carrying the R2Tg payload, a strategy intended to reduce transcriptional interference (**Fig. 3B**). We further inserted two *A. thaliana* introns^45^ into the R2Tg protein coding sequence to suppress silencing of the R2 editor system (**Fig. 3B**). Initial efforts to express R2 proteins in *N. benthamiana* showed that protein expression could be significantly improved by the introduction of geminiviral replicons leveraging autonomous replication. In line with this finding, overexpressing the complete all-in-one editor system could boost integration efficiency by increasing the overall R2 editor DNA copy number. Following these insights, we set out to test whether separating the R2Tg editor components into two constructs (RT2g-2), introducing introns into the R2Tg coding sequence (RT2g-i), and flanking the full editor system with geminiviral replicons (GV-RT2g) would improve targeted integration efficiency at the 25S rDNA site in plants (**Fig. 3B**).

Following the same method of *N. benthamiana* leaf *Agro*-infiltration and quantifying number of mCherry expressing cells from confocal images, we found that the separation of R2Tg editor components into 2 plasmids improved integration efficiency by 1.7-fold, while the introduction of introns into the R2Tg coding sequence combined with the 2-plasmid delivery approach resulted in a 2.7-fold improvement compared to the 1-plasmid delivery without introns (**Fig. 3C, E; Supplementary Fig. 3A** for more replicates). Addition of geminiviral elements to the complete 1-plasmid R2Tg editing system led to a 3.3-fold increase in integration efficiency (**Fig. 3C, E; Supplementary Fig. 3A**), achieving highest integration efficiency of ∼1.8% in this set (GV-RT2g-1). Together, these findings support the hypothesis that R2Tg protein silencing and transcriptional interference in the initial all-in-one constructs limit integration efficiency, and that strategies aimed at reducing both effects and increasing the copy number of both the R2Tg protein and payload cassettes can significantly enhance targeted integration at the 25S rDNA site.

Successful applications of R2 editors in mammalian cells consistently incubate cells transformed with editors at 37°C before detecting editing events^29–32^. The R2 editor functional screen in *N. benthamiana*, however, was conducted under standard plant growth conditions of 22°C during the light period and 20°C during the dark period in a 16-hour light/8-hour dark photoperiod. Given the high body temperature of native organisms with R2, we hypothesized that the lower temperatures used in plant studies may be limiting the catalytic efficiency of R2-mediated TPRT, and that elevating the temperature via heat shock may enhance integration activity.

To test this, *Agrobacterium tumefaciens*–infiltrated plants were first maintained at 25°C for 48 hours under a 16-hour light/8-hour dark cycle, allowing sufficient time for T-DNA delivery and accumulation of the R2 protein and RNA payload prior to the induction of heat shock. Plants were then exposed to a heat shock regimen of 25°C during the light period and 37°C during the dark period for an additional four days before analysis. Heat shock treatment of the single-plasmid R2Tg-1 system resulted in a 1.9-fold increase in targeted integration efficiency compared to the non-heat-shocked single-plasmid R2Tg-1 system. When applied to the geminiviral-amplified GV-R2Tg-1 system, heat shock yielded a 1.27-fold improvement, corresponding to a 4.2-fold increase relative to the baseline R2Tg-1 construct, reaching up to ∼2% targeted integration efficiency (**Fig. 3D, E; Supplementary Fig. 3A** for more replicates). These findings revealed that the elevated temperature indeed enhances R2Tg-mediated TPRT *in planta*.

### The R2Tg editor system achieves integration into A. thaliana and N. benthamiana protoplasts

To enable a more streamlined and quantitative workflow for assessing R2Tg-mediated integration at single-cell resolution using flow cytometry, we delivered all-in-one R2Tg editor constructs, with or without geminiviral elements, into *A. thaliana* protoplasts using polyethylene glycol (PEG)-mediated transformation. The baseline R2Tg-1 system yielded a targeted integration efficiency of 0.765%, while the geminiviral-amplified GV-R2Tg-1 system achieved 1.167% **(Supplementary Fig. 3B**), as reported by flow cytometry referring to % cells with mCherry fluorescence. Although these results demonstrate that the R2Tg system is active in *A. thaliana* protoplasts, in addition to *N. benthamiana* leaves, protoplast viability declines significantly after 48 hours, limiting the number of intact cells and window for achieving and detecting integration events.

To address this limitation, we established a complementary assay transforming *N. benthamiana* leaves via *Agro*-infiltration, and then extracting protoplasts from this transformed tissue, which allows longer incubation of the R2Tg editor system *in planta*. R2Tg-1 constructs were delivered by *Agrobacterium* infiltration, and targeted integration was detected and quantified 5 days post-infiltration in protoplasts extracted from infiltrated leaves using flow cytometry measuring mCherry fluorescence. This system yielded substantially higher integration efficiencies, reaching 4.5% with the R2Tg-1 construct **(Supplementary Fig. 3C**).

These findings confirm that the R2Tg editor system drives targeted integration into the 25S rDNA locus in both protoplast and leaf cells of two plant species. They further demonstrate that *A. thaliana* protoplasts provide a functional platform for early-stage detection, while intact leaves support extended assay durations and higher integration efficiencies for downstream applications.

### Compact RNA template with minimal homology length and 3**′** ribozyme achieves the highest integration efficiency

Molecular characterization of integration junctions at the 25S rDNA locus is critical to confirm the activity and specificity of R2Tg-mediated targeted insertion. However, the repetitive structure of rDNA tandem arrays and the presence of identical homology arms within the retrotransposition reporter generate false-positive chimeric PCR products, complicating unambiguous identification of genuine integration events^51^. In mammalian systems, such artifacts were successfully eliminated by delivering both components of the R2 editor system as RNA through the PRINT (precise RNA-mediated insertion of transgenes) method^29^. Given the technical limitations of the RNA delivery in plants, we designed a modified mCherry reporter harboring minimal homology arms of 33 base pairs upstream (R33) and 4 base pairs downstream (R4) relative to the 25S rDNA cleavage site (**Fig. 4A**). We reasoned that reducing the homology arm length would minimize PCR-generated chimera formation, thereby enabling more accurate quantification of authentic integration events without compromising efficiency, consistent with observations in mammalian cells^29,32^.

**Figure 4.**
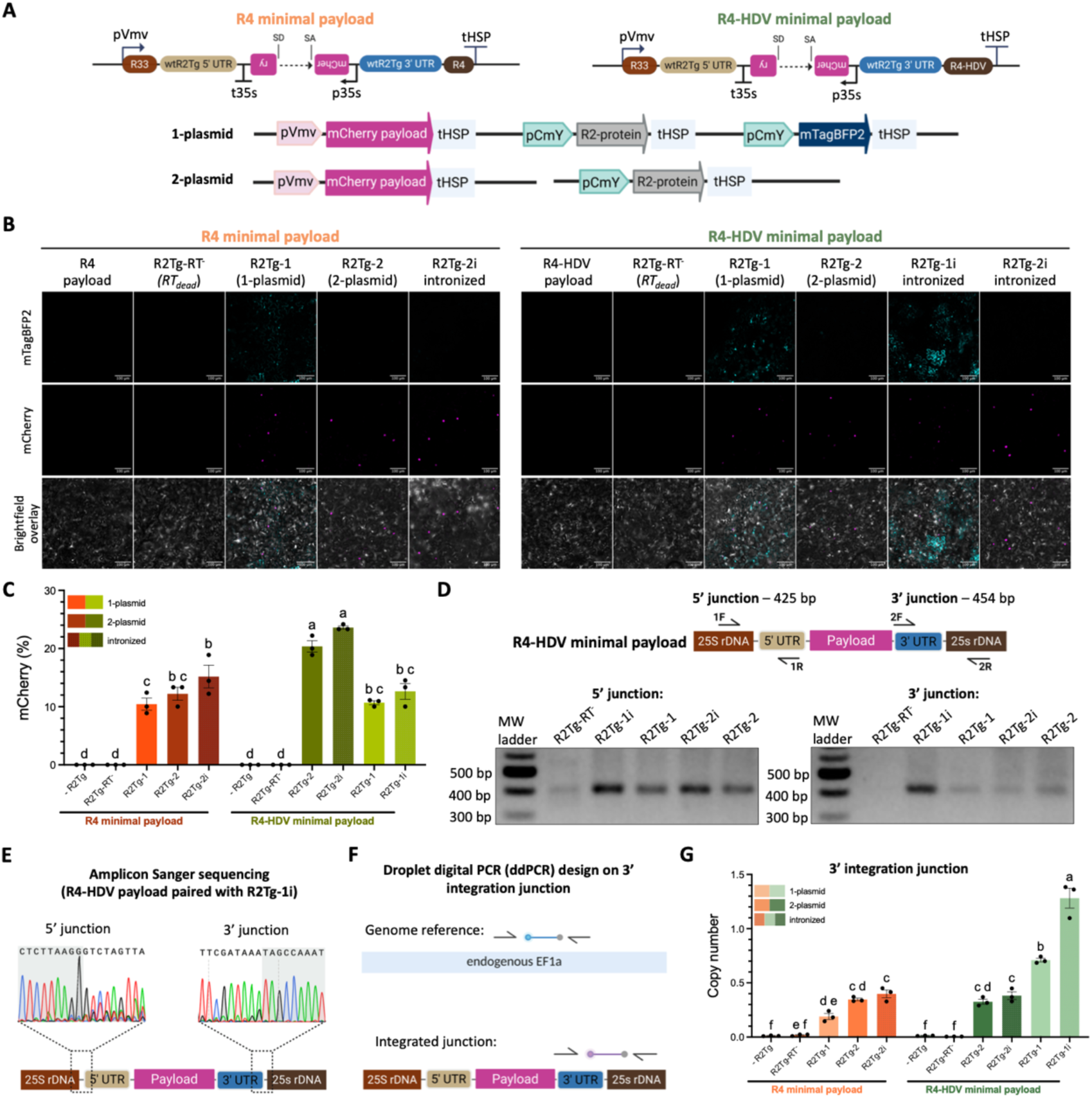
Increased integration efficiency and molecular validation via minimal homology length RNA templates with HDV ribozyme. **(A)** Schematics of R4 and R4-HDV minimal payload RNA templates and two delivery formats. **(B)** Representative confocal microscopy images of *N. benthamiana* leaves infiltrated with constructs encoding different R2Tg variants and donor RNA templates. mTagBFP2 (cyan, reporter for transformation), mCherry (magenta, R2-mediated integration reporter), and brightfield overlay. Scale bars, 100 µm. **(C)** Quantification of mCherry positive nuclei as a measure of targeted insertion efficiency. Bars represent mean ± S.E.M. of 3 independent biological replicates. P values were obtained using ordinary one-way ANOVA with Tukey HSD multiple comparison test. Different letters denote statistically significant differences. See Source Data File for details. **(D)** Target-junction PCR on genomic DNA isolated from infiltrated leaves. Expected band sizes for the 5′ and 3′ junctions are 425 bp and 454 bp. **(E)** Representative Sanger sequencing chromatograms of 5’ and 3’ integrated junction amplicons for the R4-HDV payload paired with intronized R2Tg (R2Tg-1i). **(F)** Droplet digital PCR (ddPCR) assay schematic for quantification of integration efficiency using primers and a probe specific to the integrated 3′ junction, normalized against an endogenous *NbEF1α* control locus. **(G)** Quantification of targeted integration efficiencies by ddPCR. Bars represent mean ± S.E.M. from 3 independent biological replicates. Statistical analysis is done using ordinary one-way ANOVA with Tukey HSD multiple comparison test. Different letters denote statistically significant differences. See Source Data File for details. R2Tg: *Taeniopygia guttata* R2 element, RT: reverse transcriptase, rDNA: ribosomal DNA, UTR: untranslated region, HDV: hepatitis delta virus ribozyme, R4: 4-nucleotide sequence derived from the 25S rDNA region immediately downstream of the target site, R33: 33-nucleotide sequence derived from the 25S rDNA region immediately upstream of the target site, wt: wild type, pVmv: Cassava vein mosaic virus promoter, tHSP: *Arabidopsis thaliana* heat shock protein 18.2 terminator, pCmY: pCmYLCV6 promoter derived from *Cestrum* yellow leaf-curling virus, p35s: Cauliflower mosaic virus 35S promoter, t35s: Cauliflower mosaic virus 35S terminator, SD: splice donor, SA: splice acceptor.

We also replaced the Cassava vein mosaic virus promoter (pVmv) that drives the expression of the mCherry reporter with the stronger Cauliflower mosaic virus 35S promoter (p35s) (**Fig. 4A**). Additionally, to generate a homogeneous and precise 3′ RNA end, we incorporated the hepatitis delta virus (HDV) ribozyme^52^ downstream of the payload. This ribozyme enables autocatalytic cleavage, resulting in precise processing of the RNA transcript and retention of the R4 sequence at the 3′ terminus of the R2Tg retrotransposition reporter. These modifications yielded two distinct RNA template designs, termed the R4 minimal payload and the R4-HDV minimal payload, respectively (**Fig. 4A**).

We evaluated the integration efficiencies of R2Tg editor system with the R4 and R4-HDV payload designs in *N. benthamiana* leaves. Constructs were delivered as a 1-plasmid or a 2-plasmid system separating protein and payload cassettes (**Fig. 4A**). Confocal microscopy performed 7 days post-infiltration revealed consistent mCherry signals, indicative of successful R2-mediated integration for both payload designs when paired with the wild-type R2Tg protein or an intronized version of the protein (**Fig. 4B, Supplementary Fig. 4A** for more replicates). Quantification of the percentage of cells expressing mCherry from confocal images revealed that the new designs have substantially improved the integration efficiencies. In the case of R4 minimal payload design, efficiency of R2Tg-1 increased from 0.5% to 10.4%, R2Tg-2 increased from 0.85% to 12.2%, and R2Tg-2i increased from 1.6% to 15.1%. In the case of R4-HDV minimal payload design, efficiency of R2Tg-1 reached up to 10.7%, R2Tg-1i up to 12.6%, R2Tg-2 up to 20.3%, and R2Tg-2i up to 23.6%. Notably, R2Tg-mediated integration was markedly increased in samples expressing the R4-HDV payload compared to the R4 payload (**Fig. 4B, C; Supplementary Fig. 4A**). In addition, intronized R2Tg variants delivered via either 1- or 2-plasmid strategies consistently generated the strongest mCherry signals, while the R2Tg RT-dead mutant^29^ did not produce any detectable fluorescence neither the payload without the R2 protein (**Fig. 4B, C; Supplementary Fig. 4A**).

Target-junction PCR assays corroborated imaging results at the molecular level, producing amplicons of expected sizes (425 bp for 5′ junction, 454 bp for 3′ junction) in samples harboring functional R4-HDV R2Tg minimal payload variant (**Fig. 4D**). Both 5′ and 3′ junction PCR products were detected for intronized and wild-type R2Tg constructs paired with the R4-HDV minimal payload, whereas RT-dead controls lacked the correct-sized band. Sanger sequencing junction amplicons from R4-HDV payload paired with the intronized R2Tg (R2Tg-1i) revealed precise junction sequences, perfectly matching the expected integration architecture at both 5′ and 3′ ends (**Fig. 4E**). These results validate the fidelity of the integration events generated by the optimized R2Tg editor system.

To rigorously quantify integration efficiency, we implemented a droplet digital PCR (ddPCR) assay targeting the unique 3′ and 5’ integration junctions and normalized it against the endogenous *NbEF1α* locus (**Fig. 4F; Supplementary Fig. 4B**). ddPCR analysis across 3 biological replicates showed that R2Tg editor comprising of the intronized R2Tg with the R4-HDV minimal payload in the 1-plasmid delivery format (R2Tg-1i) significantly exceeds efficiencies observed with all other R2 editor configuration tested in *N. benthamiana*, reaching an average of approximately 1.2 integrated 3′ junction copy per genome, while ∼80-90% of those copies show full-length integration events (**Fig. 4G; Supplementary Fig. 4C, D**). ddPCR at both junction sites further revealed that, across R2Tg editors tested in *N. benthamiana*, the proportion of integrated copies containing both 3′ and 5′ junctions (full-length integration events) ranged from 73% to 87% **(Supplementary Fig. 4D**).

Deep next-generation sequencing (NGS) of 3′ junction amplicons revealed 72.14% of reads harboring integrated junctions exclusively in samples expressing the R4-HDV payload in combination with intronized R2Tg (R2Tg-1i), while samples containing the payload alone lacked any integration-specific reads **(Supplementary Fig. 4E-G**). Alignment of sequencing reads confirmed high-fidelity, precise integration accompanied by a minority of indel variants comprising non-templated adenine additions and subtle shifts in endonuclease cleavage, occurring one to two nucleotides upstream of the canonical nick position, consistent with the established activity profile of R2 endonuclease **(Supplementary Fig. 4H**). Collectively, these findings demonstrate that compact R2Tg payloads designs, with minimized homology arm length and 3′ ribozyme substantially enhance R2Tg-mediated targeted integration efficiencies at the 25S rDNA site in plants and maintains precise, targeted DNA insertion.

To evaluate whether the optimized R2Tg system could mediate targeted DNA insertion at the 25s rDNA target site in the non-model crop *Solanum lycopersicum* (tomato), we tested the minimal R4-HDV mCherry payload delivered in either a one-plasmid or two-plasmid format together with the intronized R2Tg using a transient transformation method developed in our lab^53^ **(Supplementary Fig. 5A**). Confocal microscopy of seedling leaves transformed with these constructs revealed strong mCherry fluorescence in nuclei of cells co-expressing the intronized R2Tg, consistent with R2-mediated retrotransposition, while mTagBFP2 served as a transformation control **(Supplementary Fig. 5B**). PCR assays specifically targeting integrated junctions at the molecular level, further validated precise insertion of the mCherry payload at the 25S rDNA locus with the expected 425 bp and 454 bp products detected for the 5′ and 3′ junctions, respectively, while no bands were obtained when only template is delivered without the R2 protein **(Supplementary Fig. 5C-E)**. These results demonstrate that the R2Tg editor achieves efficient targeted integration in *S. lycopersicum*, establishing its functionality beyond model systems and highlighting its potential as a broadly applicable technology for precise genome engineering in crop species.

Next, we assessed whether the optimized R2Tg editor could support efficient targeted insertion of larger DNA sequences into the 25s rDNA loci of *N. benthamiana*. For this, we constructed a 5 kb R4-HDV RUBY reporter cassette^54^ (**Fig. 5A**). The RUBY reporter encoded a polycistronic open reading frame linking 3 enzymes: cytochrome P450 CYP76AD1 that oxidizes tyrosine to L-DOPA and cyclo-DOPA; L-DOPA 4,5-dioxygenase (DODA) that converts L-DOPA to betalamic acid; and a glucosyltransferase that glycosylates betanidin to produce the red betalain pigment. The three enzymes were separated by 2A peptides to allow coordinated expression from a single promoter^54,55^. The R2Tg RUBY reporter was interrupted by a plant intron in the sense orientation, similar to the mCherry payload, but with the intron placed within the glucosyltransferase gene so that betalain production occurs only upon successful R2-mediated integration of the full 5 kb polycistronic payload. The system was delivered either as a single plasmid encoding both the intronized R2Tg and the RUBY payload, or as two separate plasmids carrying R2Tg and the payload separately (**Fig. 5A**).

**Figure 5.**
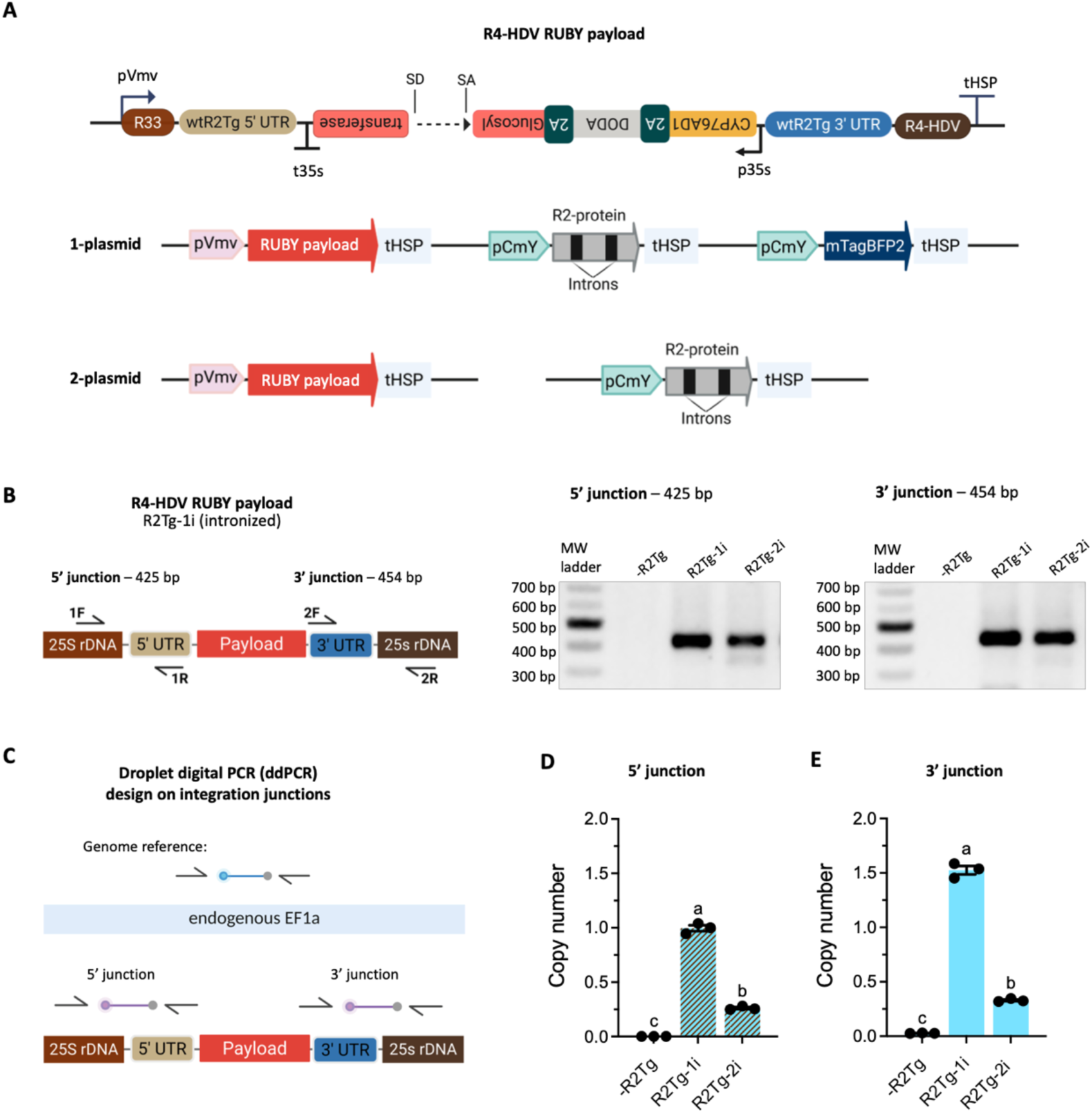
Optimized R2Tg system achieves efficient targeted insertion of a large DNA payload expressing the RUBY biosynthetic pathway. **(A)** Schematics of R4-HDV RUBY payload (5 kb) with two delivery setups. **(B)** Target-junction PCR on genomic DNA isolated from infiltrated leaves. Expected band sizes for the 5′ and 3′ junctions are 425 bp and 454 bp. **(C)** Schematic of ddPCR on the 3’ and 5’ integrated junctions. **(D)** Quantification of targeted integration efficiencies by ddPCR on the 5’ and **(E)** 3’junction. Bars represent mean ± S.E.M. from 3 independent biological replicates. Statistical analysis is done using ordinary one-way ANOVA with Tukey HSD multiple comparison test. Different letters denote statistically significant differences. See Source Data File for details. RT: reverse transcriptase, rDNA: ribosomal DNA, UTR: untranslated region, HDV: hepatitis delta virus ribozyme, R4: 4-nucleotide sequence derived from the 25S rDNA region immediately downstream of the target site, R33: 33-nucleotide sequence derived from the 25S rDNA region immediately upstream of the target site, wt: wild type, pVmv: Cassava vein mosaic virus promoter, tHSP: *Arabidopsis thaliana* heat shock protein 18.2 terminator, pCmY: pCmYLCV6 promoter, p35s: Cauliflower mosaic virus 35S promoter, t35s: Cauliflower mosaic virus 35S terminator, SD: splice donor, SA: splice acceptor. RUBY: An artificial polycistronic open reading frame that expresses the three enzymes required for Betalain biosynthesis, CYP76AD1: cytochrome P450 enzyme converts tyrosine to L-DOPA (L-3,4-dihydroxyphenylalanine), DODA: l-DOPA 4,5-dioxygenase enzyme cleaves L-DOPA to produce betalamic acid, Glucosyl transferase: glycosylates betanidin to generate the red colored betalain compound, 2A: self-cleaving 2A peptide sequence.

Expression of the RUBY payload with intronized R2Tg in *N. benthamiana* leaves produced visible betalain pigmentation within infiltrated areas, with stronger pigmentation observed for R2Tg-1i compared to R2Tg-2i, while leaves infiltrated with the RUBY payload alone, without the R2 protein, did not show pigmentation **(Supplementary Fig. 6B-D**). Target-junction PCR using the genomic DNA extracted from infiltrated *N. benthamiana* leaves confirmed successful integration, producing the expected 5’ (425 bp) and 3’ (454 bp) amplicons (**Fig. 5B**). To quantify insertion efficiencies, we applied a droplet digital PCR (ddPCR) assay specific to the integrated 5′ and 3′ junctions (**Fig. 5C**). ddPCR measurements showed that the one-plasmid system achieved the highest efficiencies, with an average of one copy per genome at the 5′ junction and 1.5 copies per genome at the 3′ junction, significantly outperforming the two-plasmid format which had 0.26 copies per genome at the 5′ junction and 0.33 copies per genome at the 3′ junction (**Fig. 5D, E**). These findings validate that the intronized R2Tg system efficiently inserts multi-kilobase metabolic pathways into the 25S rDNA safe harbor of *N. benthamiana*.

## Discussion

In this work, we established targeted DNA insertion in plants using the *Taeniopygia guttata* R2 non-LTR retroelement protein (R2Tg editor), which natively mediates site-specific integration through target-primed reverse transcription (TPRT)^27^. The R2 ribonucleoprotein complex achieves precise, multi-kilobase DNA integration by tightly coupling sequence-specific 25S rDNA recognition, bottom-strand nicking, primer hand-off, and reverse transcription of an RNA donor into cDNA^28^. Unlike CRISPR-Cas9 homology-directed repair (HDR), which depends on host DNA repair machinery to resolve a double-strand break using a donor DNA template, often resulting in low insertion efficiencies, the R2 mechanism bypasses the need for DSB induction entirely, avoiding competing end-joining pathways and the risk of large deletions or chromosomal rearrangements^15^.

R2 editors use RNA payloads in a similar manner to prime editing. In plants, prime editors are most effective for base substitutions and short insertions, and efficiency declines as insertion length and pegRNA design complexity increases. However, unlike prime editors that are constrained by insertion length, and guide RNA design complexity, R2 editors circumvent these constraints by directly synthesizing long cDNAs at the genomic target site during integration, leveraging the high processivity of their reverse transcriptase domains^27,28^. It remains to be demonstrated whether protein engineering can sufficiently increase processivity to support TPRT of even much larger payloads than the 5 kb demonstrated here. Another unique property of R2 editors is their targeting to the high-copy 25S rDNA locus, which embeds integrated genes in a specialized chromatin environment linked to stable transgene expression and minimal risk of disrupting essential host genes^56–58^.

Here, we harnessed these unique capabilities to achieve targeted insertion in *Nicotiana benthamiana* and *Arabidopsis thaliana* and *Solanum lycopersicum*. We optimized efficiencies through engineered RNA payload architectures and improved R2 protein expression systems and validated high-efficiency targeted insertion via molecular detection of integrated junctions and functional expression of fluorescent reporters. Our results illuminated key determinants of integration efficiency in plants. The highest measured mCherry positive cell counts were achieved with the intronized R2Tg protein and R4-HDV minimal payload while using the 2-plasmid delivery approach. In contrast, the highest integrated 3’ junction copy numbers detected, approaching one copy per genome, was obtained with an intronized R2Tg and R4-HDV minimal payload in a 1-plasmid delivery. Additionally, this R2Tg editor configuration achieved targeted insertion in *Solanum lycopersicum* expanding the tool’s capabilities to crop engineering. Notably, the R2Tg editor system also enabled efficient integration of a 5 kb metabolic pathway cassette, into the 25S rDNA safe-harbor locus, demonstrating that R2 retrotransposons can efficiently install complex metabolic pathways at a targeted genomic site.

The divergence in mCherry signals and ddPCR detected copy numbers per genome may reflect differences in RNA payload processing and the completeness of integrated products, as R2-mediated integration can yield incomplete insertion events^29,30,32,59^ and the fluorescence readout from the mCherry retrotransposition reporter requires successful intron splicing before integration. Incomplete integration events or unspliced integration events would not yield detectable mCherry signal but molecular-level detection copy number allows us to obtain a comprehensive picture of integration events at the 25S rDNA site. These observations are consistent with the possibility that R2Tg silencing and transcriptional interference in the all-in-one setup may affect efficient splicing and integration of the mCherry reporter. Intronization of R2Tg and separation of protein and payload cassettes may alleviate these constraints, improve the expression and splicing of payloads prior to integration at the 25S rDNA site to yield detectable mCherry reporter expression.

We observed improved mCherry signal after heat shock treatment of *N. benthamiana* plants, expressing the R2 editor system after *Agrobacterium*-mediated leaf infiltrated. This improvement in integration efficiency may be a result enzymatic kinetics or ribonucleoprotein (RNP) interactions required for integration. Other temperature-sensitive host processes may also contribute to this effect. In particular, heat stress has been shown to activate endogenous transposable elements in plants through global chromatin relaxation, reduction of DNA methylation, and suppression of RNA interference^46–48,50,60^. Such mechanisms may alleviate epigenetic repression of R2 editor components, thereby facilitating higher integration efficiencies. Disentangling these direct catalytic effects from host epigenetic changes will be important for understanding the mechanism at the molecular level.

In future studies, it will be important to assess whether transcriptional regulation of R2 protein and RNA template expression impose plant-specific constraints on R2 integration efficiencies not seen mammalian systems. To complement our work further, comprehensive genome-wide mapping is required to confirm the absence of off-target integrations, and long-term studies of stability and heritability will be crucial for translating this platform into breeding and crop improvement pipelines.

Although in this study we delivered the R2 system using *Agrobacterium tumefaciens*, which introduces DNA randomly into the genome, future efforts will focus on direct delivery of the R2 system as a ribonucleoprotein complex using nanoparticles or biolistics. This strategy would eliminate unintended T-DNA insertions, ensuring that only precise edits at the target rDNA locus are recovered. Such delivery advances will not only improve editing fidelity but also accelerate generation of clean, marker-free lines, reduce the need for laborious screening, and broaden applicability across plant species that are recalcitrant to *Agrobacterium*. Together, these improvements could establish R2-based editors as a practical and widely deployable platform for precise, large-fragment DNA insertion in plants, an advancement with far-reaching implications for both fundamental genome studies and crop engineering.

Together, these findings establish R2 retrotransposons as a distinctive and versatile addition to the plant genome engineering toolkit. By leveraging the precision of a native retroelement and augmenting it through improved protein expression platforms and engineered RNA payloads designs, this technology enables efficient, DSB-free integration of gene-sized cassettes into a defined genomic safe harbor locus in plants. With continued refinement, R2-based systems have the potential to complement and, in certain applications, surpass existing technologies for precise and targeted introduction of complex genetic traits into plant genomes.

## Methods

### Plasmid construction

Coding sequences for 4 R2 non-LTR retrotransposases, R2Tg (*Taeniopygia guttata*), R2Bm (*Bombyx mori*), R2Tg-opt (rationally engineered *T. guttata*) and R2Za (*Zonotrichia albicollis*), were codon-optimized for *Nicotiana benthamiana* and *Arabidopsis thaliana* using GenSmart Codon Optimization (GenScript) and synthesized as double-stranded DNA fragments (Twist Bioscience). Each open reading frame contained an N-terminus bipartite SV40 nuclear localization signal (KRTADGSEFESPKKKRKV). 3 additional versions of the R2Tg protein were generated: i) R2Tg-intronic, which contains 2 *A. thaliana* introns^45^; ii) R2Tg RT-dead, carrying D660A and D661A substitutions in the reverse-transcriptase domain; and iii) R2Tg EN-dead, carrying D1057A and D1070A substitutions in the endonuclease domain^29^.

The retrotransposition reporter followed a classic GFP-intron design^42,43^. A Cauliflower mosaic virus (CaMV) 35S promoter and an *A. thaliana* HSP18.2 terminator flank a reverse-oriented nested cassette that contains the Cassava vein mosaic virus (CsVMV) promoter, mCherry open reading frame interrupted by either the *Solanum tuberosum* ST-LS1 intron or an *A. thaliana* intron, and a CaMV t35S terminator^44,45^. Plasmids carrying R2 protein expression cassettes and donor DNA templates were combined with Loop Assembly using BsaI- and SapI-mediated Golden Gate reactions (NEB catalog numbers R0569S and E1601L respectively) and cloned into pCAMBIA backbones^61^. All plasmids were propagated in NEB Turbo *Escherichia coli* competent cells (NEB catalog no. C2984I) and sequence-verified by Plasmidsaurus. A geminiviral replicon based on the Bean yellow dwarf virus (BeYDV) was used to transiently overexpress either the R2 proteins or the complete R2 editing system^41^. Constructs generated in this work can be found in Supplementary Table 1.

### Plant growth and transformation

*Nicotiana benthamiana* and *Arabidopsis thaliana* (Col-0) plants were grown in a Conviron controlled-environment chamber set to 22 °C during the day, 20 °C during the night with a 16 h light / 8 h dark photoperiod and 55 % relative humidity. 4-weeks old *N. benthamiana* plants were used for leaf infiltrations for transient experiments. 4-weeks old *A. thaliana* plant leaves were harvested for protoplast isolation.

*Agrobacterium tumefaciens* strain GV3101 (GOLDBIO catalogue no. CC-125) carrying the binary R2 construct, the p19 silencing-suppressor plasmid, and the pSOUP helper plasmid was cultured overnight at 28°C in 2YT medium (Invitrogen catalogue no. 12-780-052) supplemented with the appropriate antibiotics. Cells were collected and resuspended in induction media (10 mM MES pH 5.5, 10 mM MgCl_2_, 200 μM Acetosyringone (PhytoTech Labs catalogue no. A104-5G) in DMSO), at OD_600_ 0.4. Suspensions were allowed to shake at 70 rpm for 4 hours at room temperature. The *Agrobacterium* suspension was infiltrated into the abaxial surface of fully expanded leaves using a 1 mL needleless syringe until the tissue was saturated. 3 leaves per plant were treated. Plants were returned to the growth chamber immediately and sampled 3 to 7 days post infiltration depending on the experiment.

### Transient Transformation of *Solanum lycopersicum* (cv. Micro-Tom)

Transient transformation was performed using a protocol adapted from the VAST (Vacuum and Sonication-Assisted Transient expression) method^53^. Seeds of *S. lycopersicum* (cv. Micro-Tom) were surface-sterilized with 33% (v/v) bleach containing 0.1% Tween-20 for 6 min, rinsed 4–5 times with sterile water, and germinated on ½ MS agar at 22 °C under a 16 h light/8 h dark photoperiod until cotyledons were fully expanded (∼9 days). *Agrobacterium tumefaciens* strain GV3101 (GOLDBIO, CC-125) carrying the binary R2 construct, p19 silencing suppressor, and pSOUP helper plasmid was cultured in 2X-YT medium with antibiotics at 28 °C, 200 rpm, 2 days prior to co-cultivation. Cultures were pelleted (8,000 g, 5 min), resuspended in AB-MES induction medium (200 µM Acetosyringone, no antibiotics) to OD₆₀₀ = 0.3, and incubated overnight at 28 °C, 200 rpm. On the day of co-cultivation, cultures were pelleted again and resuspended in co-culture medium (1:1 AB-MES and ½ MS supplemented with 200 µM Acetosyringone) and adjusted to OD₆₀₀ = 0.3.

For infection, 7–8 seedlings were immersed in 8 mL of *Agrobacterium* suspension in 12 mL tubes, sonicated for 30 s (Branson CPX-952-116R), and subjected to three cycles of vacuum infiltration (5 min vacuum followed by rapid release per cycle) with gentle mixing between cycles. Seedlings were then transferred to six-well plates containing 4 mL of fresh *Agrobacterium* suspension per well and co-cultivated for 2 days at 22 °C under a 16 h light/8 h dark cycle. Following co-cultivation, seedlings were washed three times with sterile water and transferred to ½ MS agar plates containing 100 µM timentin to suppress bacterial growth. Plants were maintained under standard growth conditions, and transient expression was assessed 3 days post-infection by confocal microscopy.

### Confocal microscopy

Leaf discs ∼5 mm in diameter were collected from infiltrated *Nicotiana benthamiana* leaves using a sterile hole puncher, mounted in distilled water, and imaged immediately on a Leica Stellaris 8 confocal laser-scanning microscope (HC PL APO 20×/0.75 NA water-immersion objective) housed in the Biological Imaging Facility (BIF) at Caltech. The blue fluorescent protein (BFP) reporter was excited at 405 nm and its emission collected from 430 nm to 470 nm, while mCherry was excited at 561 nm and its emission collected from 580 nm to 630 nm. A single optical section was acquired for each field with the pinhole set to 1 Airy unit, using identical laser power, detector gain, and offset for all samples within an experiment. Images were exported from Leica LAS X v4.3 as 16-bit TIFF files. A 100 µm scale bar was added to exported images using ImageJ Fiji.

### Protein expression and detection

R2Tg, R2Bm, R2Tg-opt, and R2Za coding sequences were fused to an N-terminal YPet tag for detection and confirmation of nuclear localization via confocal microscopy. Complementarily, R2 proteins were fused to an N-terminal HiBiT peptide tag for higher throughput luminescence detection using the Nano-Glo® HiBiT Lytic Detection System. 3 days post infiltration (dpi), 3 infiltrated *Nicotiana benthamiana* leaves were harvested, flash-frozen in liquid nitrogen, and ground to a fine powder before extraction in lysis buffer (20 mM Tris-HCl pH 7.4 (ThermoFisher Scientific catalogue no. 15567027), 25 % glycerol (Sigma-Aldrich catalogue no. G5516), 20 mM KCl (Sigma-Aldrich catalogue no. P5405), 2 mM EDTA (Sigma-Aldrich catalogue no. E9884), 2.5 mM MgCl₂ (Sigma-Aldrich catalogue no. M2393), 250 mM sucrose (ThermoFisher scientific catalogue no. J65148.A1), 0.1 % PMSF (ThermoFisher Scientific catalogue no. 36978)). After centrifugation at 1500 × g for 10 min at 4 °C to pellet nuclei, the pellet was washed up to 5 times with wash buffer (20 mM Tris-HCl pH 7.4 (ThermoFisher Scientific catalogue no. 15567027), 25 % glycerol (Sigma-Aldrich catalogue no. G5516), 2.5 mM MgCl₂ (Sigma-Aldrich catalogue no. M2393), 0.2 % Triton X-100 (Sigma-Aldrich catalogue no. S5886)) and lysed in high-salt buffer (20 mM HEPES-KOH pH 7.9 (Sigma-Aldrich catalogue no. H3375), 2.5 mM MgCl₂ (Sigma-Aldrich catalogue no. M2393), 100 mM NaCl (Sigma-Aldrich catalogue no. S5886), 20 % glycerol (Sigma-Aldrich catalogue no. G5516), 0.2 mM EDTA (Sigma-Aldrich catalogue no. E9884), 0.5 mM DTT (GOLDBIO catalogue no. DTT), one cOmplete Mini EDTA-free protease-inhibitor tablet (Sigma-Aldrich catalogue no. 11836170001) per 10 mL). For HiBiT detection, 30 µL of nuclear lysate was combined with 30 µL Nano-Glo HiBiT Lytic reagent (Promega catalogue no. N3030) containing LgBiT and furimazine, and luminescence was recorded on a Tecan Spark plate reader with a 1000 millisecond integration time.

### Isolation and transfection of *Nicotiana benthamiana* and *Arabidopsis thaliana* protoplasts

Mesophyll protoplasts were isolated from fully expanded leaves of 4-week-old *N. benthamiana* or *A. thaliana* (Col-0) following Yoo et al. (2007)^62^ with minor modifications. On the day of the experiment, a fresh enzyme solution (20 mL) containing 20 mM MES pH 5.7 (Sigma-Aldrich catalogue no. M2933), 0.4 M mannitol (Millipore Sigma catalogue no. 443907), 20 mM KCl (Sigma-Aldrich catalogue no. P5405), 1.5% (w⁄v) cellulase onozuka R-10 (Yakult Pharmaceutical IND. CO., LTD) and 0.4% (w⁄v) macerozyme R-10 (Yakult Pharmaceutical IND. CO., LTD) was prepared; the MES–mannitol–KCl base was pre-heated to 70°C for 4 min before the enzymes were dissolved. After cooling, 10 mM CaCl₂ (Sigma-Aldrich catalogue no. C7902) and 0.1% (w⁄v) BSA (Fisher bioreagents catalogue no. BP9703) were added, and the solution was passed through a 0.45 µm syringe filter into a Petri dish. *N. benthamiana* leaf strips or *A. thaliana* full leaves were incubated in this solution without agitation for 3 h at room temperature in the dark, with gentle swirling once per hour. The digest was diluted 1:1 with W5 solution (154 mM NaCl (Sigma-Aldrich catalogue no. S5886), 125 mM CaCl₂ (Sigma-Aldrich catalogue no. C7902), 5 mM KCl (Sigma-Aldrich catalogue no. P5405), 2 mM MES pH 5.7 (Sigma-Aldrich catalogue no. M2933)), passed through a 70 µm nylon mesh (Fisherbrand catalogue no. 22363548), and the flow-through was centrifuged at 200 × g for 2 min. The pellet was gently resuspended in 5 mL W5, chilled on ice for 30 min, then the supernatant was removed and the protoplasts were adjusted to 5 × 10⁵ cells mL⁻¹ in MMG solution (0.4 M mannitol (Millipore Sigma catalogue no. 443907), 15 mM MgCl₂ (Sigma-Aldrich catalogue no. M2393), 4 mM MES pH 5.7 (Sigma-Aldrich catalogue no. M2933)).

For PEG-mediated transfection a 40 % PEG 4000 solution (Sigma-Aldrich catalogue no. 81240) was freshly prepared (4 g PEG 4000, 3 mL H₂O, 2.5 mL 0.8 M mannitol (Millipore Sigma catalogue no. 443907), 1 mL 1 M CaCl₂ (Sigma-Aldrich catalogue no. C7902)). In round-bottom Eppendorf tubes, 20 µg plasmid DNA (≤ 20 µL) was combined with 100 µL protoplast suspension, mixed gently, and 120 µL PEG solution was added. After 15 min at room temperature, 1 mL W5 was added and the mixture was centrifuged (50 × g, 2 min); the wash was repeated twice. The pellet was resuspended in WI solution (0.5 M mannitol (Millipore Sigma catalogue no. 443907), 20 mM KCl (Sigma-Aldrich catalogue no. P5405), 4 mM MES pH 5.7 (Sigma-Aldrich catalogue no. M2933)), transferred to BSA-blocked well plates (0.1 % BSA (Fisher bioreagents catalogue no. BP9703)), and incubated at room temperature in the dark for 24-72 h before downstream analyses._After incubation in the dark at room temperature for 24 to 72 h, transformed protoplasts were used for confocal imaging and/or flow cytometry. In a modified protocol, protoplasts were extracted from 5-week-old *Nicotiana benthamiana* leaves 7 days after *Agrobacterium* infiltration and analyzed directly by flow cytometry.

### Flow cytometry analysis

Protoplasts were analyzed on a CytoFLEX flow cytometer (Beckman Coulter) provided by the Caltech Flow Cytometry and Cell Sorting Facility. Live protoplast gates were defined using fluorescein diacetate (FDA) staining (Fisher scientific catalogue no. F1303), detected in the FITC channel. BFP was excited with a 405 nm laser and detected in the 450/45 nm channel, while mCherry was excited with a 561 nm laser and detected in the 610/20 nm channel. Data was exported as FCS files and analyzed using FloReada for gating, quantification, and visualization.

### Amplicon PCR and sanger sequencing

Genomic DNA (gDNA) was extracted from *Nicotiana benthamiana* leaf tissue infiltrated with R2 constructs using the DNeasy Plant Mini Kit (Qiagen catalogue no. 69106) according to the manufacturer’s protocol. For 3′-junction analysis, 25 µL PCRs were assembled with 1× Phusion High-Fidelity Master Mix (NEB catalogue no. M0531S), 0.5 µM each primer 2F and 2R, and 20 ng gDNA. Cycling conditions: 98 °C for 30 s; 35 cycles of 98 °C for 10 s, 60 °C for 20 s, 72 °C for 30 s; final extension 72 °C for 2 min. The expected 450 bp amplicons were verified on a 1 % agarose gel, purified with the QIAquick Gel Extraction Kit (Qiagen catalogue no. 28704) and submitted to Laragen Inc. for Sanger sequencing. Chromatograms were aligned to reference sequences in SnapGene v6.2 to confirm precise junction formation.

### Next generation sequencing (NGS)

The 450 bp 3′-junction PCR products (primer pair 2F/2R) were gel-purified with the QIAquick Gel Extraction Kit (Qiagen catalogue no. 28704) and eluted in 35 µL nuclease-free water. DNA concentration was measured on a NanoDrop 2000 spectrophotometer, and each amplicon was adjusted to 50 ng/µL, sealed in Eppendorf microcentrifuge tubes, and shipped overnight to the Massachusetts General Hospital CCIB DNA Core (Harvard) for Illumina sequencing. The core performed NGS and provided (i) de novo-assembled contigs representing variants present at > 1 % abundance and (ii) the complete demultiplexed FASTQ files.

In Galaxy, paired-end reads were processed with DADA2: adapters were trimmed, reads were quality-filtered, and overlapping pairs were merged. Merged reads were aligned to a custom reference comprising the payload integrated at the 25S rDNA locus using Bowtie2 v2.5.0 to capture low-frequency variants (< 1%). The subset of sequence variants mapping perfectly to the insertion junction were imported into Geneious Prime 2024.1, aligned to the same reference and the multiple alignment was exported as FASTA. Alignment visualization was carried out in Google Collab with Biopython 1.81, pandas 2.2.2, and matplotlib 3.7.3; the resulting grid plot of base-by-base variation was exported as high-resolution PNG files for publication.

### Droplet digital PCR (ddPCR) for detecting targeted insertion efficiency

Genomic DNA was extracted from 100 mg of *Nicotiana benthamiana* leaf tissue collected from the *Agrobacterium* infiltration site 7 days after infiltration using the DNeasy Plant Mini Kit (Qiagen). Genomic DNA was predigested with FastDigest KpnI (Thermo Fisher Scientific, catalog no. FD0524) according to the manufacturer’s protocol. Duplex 22 µl ddPCR reactions were prepared by mixing 11 µl of ddPCR Supermix for Probes (no dUTP; Bio-Rad, catalog no. 1863024), 900 nM forward and reverse primers for the target and reference loci, 250 nM FAM-labeled probe targeting the *NbEF1α* reference gene, 250 nM HEX-labeled probe targeting the integrated R2 junction, and 50 ng of predigested gDNA. Oligonucleotide sequences were synthesized by Integrated DNA Technologies (IDT) and are listed in Supplementary Table 2. Reaction mixtures were loaded into a DG8 cartridge (Bio-Rad, catalog no. 1864007) along with 70 µl of droplet generation oil (Bio-Rad, catalog no. 1863005), and droplets were generated using a Bio-Rad QX200 Droplet Generator. Following droplet generation, 40 µl of emulsion was transferred to a 96-well plate, heat-sealed with pierceable foil, and thermal-cycled under the manufacturer’s conditions with an annealing/extension temperature of 55.° C for 3’ junction and 60°C for 5’ junction. Droplets were read using a Bio-Rad QX200 Droplet Reader. Data were analyzed using QX Manager software (Bio-Rad). All Bio-Rad ddPCR equipment was provided by the CLARITY, Optogenetics and Vector Engineering Research (CLOVER) Center at Caltech. Insertion efficiency was calculated as the ratio of HEX-labeled target copy number to the FAM-labeled reference gene (*NbEF1α*) copy number assuming 4 copies of the target sequence.

## Data Availability Statement

All data are included either in the manuscript or supplementary files.

## Author Contributions

Conceptualization: KTM, GSD

Methodology: KTM, YW, TMO, AS, EL, GSD

Data Acquisition: KTM, YW, TMO, AS, EL

Supervision: KTM, TMO, GSD

Writing: KTM, GSD

Funding Acquisition: GSD

## Acknowledgements

We thank Professor Kathleen Collins (University of California, Berkeley) for valuable discussions and insights on experimental design. Imaging was performed in the Biological Imaging Facility, with the support of the Caltech Beckman Institute and the Arnold and Mabel Beckman Foundation. We acknowledge the Beckman Institute CLOVER Center for providing access to droplet digital PCR instrumentation and support with experiments. Flow cytometry data used in this manuscript was generated in the Caltech Flow Cytometry & Cell Sorting Facility (Research Resource Identifier number is RRID: SCR_025087). Schematics were created with BioRender.com.

