## Supplementary Figures for "Optimized R2 Retroelement Complexes Enable Precise and Efficient DNA Insertion into Plant Genomes": R2 supp figures.docx

**Supplementary Figure 1. Validation of 5′ UTR autocleavage activity and functional screening of R2 editors. (A)** Schematic of the *in vitro* 5′ UTR autocleavage assay. 25s rDNA 100 bp homology sequence was fused to R2 5′ untranslated regions (5′ UTRs) downstream of a T7 promoter, which was transcribed *in vitro* and assayed for autocleavage activity based on fragment sizes. **(B)** 2% TAE agarose gel showing cleavage products of three 5′ UTR variants: minimal *Tribolium castaneum* (minR2Tc), wild-type *T. castaneum* (wtR2Tc), and wild-type *Taeniopygia guttata* (wtR2Tg). Orange, blue and grey arrowheads indicate expected fragments. Band sizes match predicted cleavage outcomes: minR2Tc (uncleaved~177 nt, 105 nt, 72 nt), wtR2Tc (uncleaved~265 nt, 193 nt, 72 nt), and wtR2Tg (uncleaved~275 nt, 203 nt, 72 nt). A 300 nt synthetic RNA template was used to denote uncleaved template sizes. **(C)** Representative confocal microscopy images from the R2 editor function screen in *N. benthamiana* leaves, displaying mTagBFP2 (cyan, delivery marker) and mCherry (magenta, reporter of successful integration) fluorescence for four R2 protein variants: R2Bm (*Bombyx mori*), R2Tg (*T. guttata*), codon-optimized R2Tg-OPT, and R2Za (*Zonotrichia albicollis*) combined with four donor RNA architectures that pair distinct 5′/3′-UTR sets: Green: minR2Tc 5′ UTR + wtR2Gf 3′ UTR (minimal *Tribolium castaneum* 5′ UTR with wild-type *Geospiza fortis* 3′ UTR); Yellow: wtR2Tc 5′ UTR + wtR2Gf 3′ UTR (wild-type *T. castaneum* 5′ UTR with wild-type *G. fortis* 3′ UTR); Purple: wtR2Tg 5′ UTR + wtR2Tg 3′ UTR (wild-type *T. guttata* UTRs); and Black: minR2Tg 5′ UTR + minR2Tg 3′ UTR (wild-type *T. guttata* 5’ UTR and minimal *T. guttata* 3’ UTR). Scale bars, 100 µm. White arrow heads indicate nuclear mCherry signal coming from the R2-mediated targeted integration. LHA: left homology arm, UTR: untranslated region.

**Supplementary Figure 2. Negative control samples from the functional screening of R2 editors. (A)** Schematic of the R2 editor function screen showing four R2 protein variants, R2Bm (*Bombyx mori*), R2Tg (*Taeniopygia guttata*), R2Tg-OPT (a rationally improved variant of R2Tg), and R2Za (*Zonotrichia albicollis*). R2 variants are paired with different 5′ and 3′ untranslated region (UTR) combinations with distinct 5′/3′-UTR sets: Green: minR2Tc 5′ UTR + wtR2Gf 3′ UTR (minimal *Tribolium castaneum* 5′ UTR with wild-type *Geospiza fortis* 3′ UTR); Yellow: wtR2Tc 5′ UTR + wtR2Gf 3′ UTR (wild-type *T. castaneum* 5′ UTR with wild-type *G. fortis* 3′ UTR); Purple: wtR2Tg 5′ UTR + wtR2Tg 3′ UTR (wild-type *T. guttata* UTRs); and Black: minR2Tg 5′ UTR + minR2Tg 3′ UTR (wild-type *T. guttata* 5’ UTR and minimal *T. guttata* 3’ UTR). All R2 payload cassettes used in the R2 editor function screen carry 100-bp left and right homology arms. **(B)** Representative confocal fluorescence microscopy images of *N. benthamiana* leaves infiltrated with constructs encoding only the payload cassettes and lacking the R2 protein effector. mTagBFP2 (cyan) reports transformation, mCherry (magenta) would report R2-mediated integration in the presence of an R2 protein effector. Scale bars, 100 µm. UTR: untranslated region.

**Supplementary Figure 3. Confocal imaging and flow cytometry analysis of R2Tg and GV-R2Tg in leaves and protoplasts.** **(A)** Representative confocal fluorescence images of *N. benthamiana* leaves infiltrated with R2Tg constructs that express the R2Tg protein and donor template from a single plasmid, from separate plasmids, or an intronized R2Tg expression cassette, together with GV-R2Tg constructs. mTagBFP2 (cyan) marks construct delivery, and mCherry (magenta) reports payload integration and expression. Two biological replicates are shown for each condition. Scale bars, 100 µm. **(B)** Flow-cytometry dot plots of *Arabidopsis thaliana* protoplasts transiently transfected with R2Tg-1 plasmid or GV-R2Tg-1 plasmid constructs, displaying mTagBFP2 and mCherry fluorescence. **(C)** Schematic of *N. benthamiana* leaf infiltration and protoplast isolation workflow. Flow cytometry plots of *N. benthamiana* protoplasts showing mTagBFP2 and mCherry fluorescence following infiltration with mTagBFP2 and mCherry positive control samples, and R2Tg-1 plasmid constructs. BFP⁺: blue fluorescent protein-positive population, mCherry⁺: red fluorescent protein-positive population, BFP⁺/mCherry⁺: dual-positive population indicating successful delivery and integration, GV: geminiviral replicon, R2Tg: *Taeniopygia guttata* R2 element, R2Tg-opt: rationally improved *T. guttata* R2 element, R2Tg-1: single-plasmid R2Tg system, R2Tg-2: dual-plasmid R2Tg system, R2Tg-2i: dual-plasmid intronized R2Tg system, GV-R2Tg-1: geminiviral single-plasmid R2Tg system, FL1-A: channel detecting BFP fluorescence (excitation 405 nm, emission ~450 nm), FL5-A: channel detecting fluorescein diacetate (FDA) signal for cell viability (excitation 488 nm, emission ~530 nm). FL8-A: channel detecting mCherry fluorescence (excitation 561 nm, emission ~610 nm).

**Supplementary Figure 4. Characterization of R4 and R4-HDV minimal payloads in *N. benthamiana* by ddPCR and NGS. (A)** Confocal fluorescence microscopy of *N. benthamiana* leaves co-infiltrated with R2Tg and either minimal R4 or R4-HDV payload constructs. mTagBFP2 (cyan) marks delivery; mCherry (magenta) indicates reporter activation. Scale bars, 100 µm. **(B)** Schematic of droplet digital PCR (ddPCR) assay for quantification of integration efficiency using primers and a probe specific to the integrated 5′ junction, normalized against an endogenous *NbEF1α* control locus. **(C)** Quantification of targeted integration efficiencies by ddPCR. Bars represent mean ± S.E.M. from 3 independent biological replicates. Statistical analysis is done using ordinary one-way ANOVA with Tukey HSD multiple comparison test. Different letters denote statistically significant differences. See Source Data File for p-values and statistical significance details. **(D)** Quantification of full-length integrated products with intact 3’ and 5’ junctions detected by ddPCR. Bars represent mean ± S.E.M. from 3 independent biological replicates. Statistical analysis is done using ordinary one-way ANOVA with Tukey HSD multiple comparison test. Different letters denote statistically significant differences. See Source Data File for p-values and statistical significance details. **(E)** Schematic of the 3′ junction showing the R2 payload flanked by 5′ and 3′ UTRs inserted into the 25S rDNA locus. Primer 2F anneals within the 3′ UTR and primer 2R anneals downstream in native 25S rDNA. **(F)** Schematic of R2 target site in the 25s rDNA locus showing nucleotides derived from the R2 3′ UTR (blue) and 25S rDNA target (brown) red arrowheads mark the inferred bottom strand nick positions for the R2 protein. **(G)** Percentage of sequencing reads containing integrated junctions in samples with the R4-HDV payload and R4-HDV paired with R2Tg-1i construct. **(H)** Representative amplicon sequencing reads aligned to the reference 3′ junction show the indel profile. Each row corresponds to a unique read, with nucleotide identity indicated by color (A: green, T: purple, C: blue, G: orange). Red boxes mark extra bases at the 3′ junction, which arise from R2 endonuclease nicking 1–2 bases upstream of the canonical integration site and/or non-templated addition of A nucleotides prior to the start of the 3′ UTR. Read counts and percentages are shown to the right. R2Tg: *Taeniopygia guttata* R2 element, rDNA: Ribosomal DNA, UTR: Untranslated region, HDV: Hepatitis Delta Virus ribozyme, R4: 4-nucleotide sequence derived from the 25S rDNA region immediately downstream of the target site, RT: Reverse transcriptase, NGS: Next-generation sequencing.

**Supplementary Figure 5. Optimized R2Tg editor system achieves targeted insertion in *Solanum lycopersicum.* (A)** Schematic of R4-HDV ribozyme minimal payload RNA template designed to report on R2Tg-mediated retrotransposition. Two delivery formats were tested: a 1-plasmid system encoding both the R2Tg (intronized) protein and payload cassette, and a 2-plasmid system with R2Tg (intronized) protein and payload cassettes delivered on separate *Agrobacterium* T-DNA. **(B)** Representative confocal fluorescence microscopy images of *S. lycopersicum* seedling leaves transformed with constructs encoding the intronized R2Tg and R4-HDV mCherry payload RNA template, imaged 3 days post-transformation. Each row represents a fluorescence channel: mTagBFP2 (cyan, reporter for transformation), mCherry (magenta, R2-mediated integration reporter), and brightfield overlay. Scale bars, 100 µm. White arrows indicate nuclear localized mCherry signal. **(C)** Schematic of 5’ and 3’ integrated junction PCRs detecting targeted insertion at the 25s rDNA target site. **(D)** 5’ and **(E)** 3’ junction PCR detects R2Tg-mediated integration. RT: reverse transcriptase, rDNA: ribosomal DNA, UTR: untranslated region, HDV: hepatitis delta virus ribozyme, R4: 4-nucleotide and R33: 33-nucleotide sequence derived from the 25S rDNA region immediately downstream of the target site, wt: wild type, pVmv: Cassava vein mosaic virus promoter, tHSP: *Arabidopsis thaliana* heat shock protein 18.2 terminator, pCmY: pCmYLCV6 promoter, p35s: Cauliflower mosaic virus 35S promoter, t35s: Cauliflower mosaic virus 35S terminator, SD: splice donor, SA: splice acceptor.

**Supplementary Figure 6. *Nicotiana benthamiana* leaves expressing the RUBY metabolic pathway produce betalain. (A)** Schematic of the 5 kb R4-HDV RUBY payload designed to report R2Tg-mediated retrotransposition. Two delivery formats were tested: a single plasmid encoding both intronized R2Tg and the RUBY cassette, or two plasmids carrying the R2Tg protein and RUBY cassette separately. **(B)** Infiltrated leaf area expressing the RUBY payload without R2Tg as a negative control. **(C)** Infiltrated leaf area expressing the RUBY payload with intronized R2Tg in the 2-plasmid format. **(D)** Infiltrated leaf area expressing the RUBY payload with intronized R2Tg in the 1-plasmid format. Scale bar, 1cm. Dashed circles mark infiltration zones. Images were taken with an iPhone 12 mini. R2Tg: *Taeniopygia guttata* R2 element, RT: reverse transcriptase, rDNA: ribosomal DNA, UTR: untranslated region, HDV: hepatitis delta virus ribozyme, R4: 4-nucleotide sequence derived from the 25S rDNA region immediately downstream of the target site, R33: 33-nucleotide sequence derived from the 25S rDNA region immediately upstream of the target site, wt: wild type, pVmv: Cassava vein mosaic virus promoter, tHSP: *Arabidopsis thaliana* heat shock protein 18.2 terminator, pCmY: pCmYLCV6 promoter derived from *Cestrum* yellow leaf-curling virus, p35s: Cauliflower mosaic virus 35S promoter, t35s: Cauliflower mosaic virus 35S terminator, SD: splice donor, SA: splice acceptor. Ruby: An artificial polycistronic open reading frame that expresses all the three enzymes required for Betalain biosynthesis, CYP76AD1: cytochrome P450 enzyme converts tyrosine to L-DOPA (L-3,4-dihydroxyphenylalanine), DODA: l-DOPA 4,5-dioxygenase enzyme cleaves L-DOPA to produce betalamic acid, Glucosyl transferase: glycosylates betanidin to generate the red colored betalain compound, 2A: self-cleaving 2A peptide sequence that enables polycistronic expression of multiple proteins from a single transcript
